# TigerAI: An AI-powered genetic evidence platform to support clinical development

**DOI:** 10.64898/2026.08.29.748046

**Authors:** Zichen Zhang, Wanheng Zhang, Xiaochen Yang, Yujue Li, Jinjie Lin, Chong Wu, Bingxin Zhao

## Abstract

Genetic evidence is a major determinant of clinical success in drug development, yet its aggregation has long relied on laborious human curation. Large language models (LLMs) have the potential to rapidly synthesize knowledge across biomedical resources, providing a route to scalable AI-driven genetic evidence generation. Here we develop a novel domain-grounded instruction framework to systematically evaluate GPT-5 for producing genetic evidence relevant to clinical trial success. Using 13,022 target-indication pairs from a comprehensive drug development database, we benchmark LLM-derived evidence against a recent exhaustive human expert-curated study. We find that GPT-5 yields genetic evidence that is at least as informative as expert curation for inferring clinical success, while substantially expanding coverage relative to traditional curation resources. Building on these results, we introduce TigerAI (https://tigerai.bio/), a dual-purpose platform for AI-powered genetic evidence that (i) benchmarks emerging state-of-the-art LLMs and (ii) provides an accessible service for querying reliable AI-generated genetic evidence. These contributions outline a practical, domain-grounded pathway for integrating AI-powered genetic evidence into drug development pipelines and for realizing the potential of LLMs to inform clinical success.

## Main

Human genetic evidence is a robust, well-established scientific determinant of clinical success in drug development^1–4^, a process that is both costly and time-consuming^5–7^. However, the generation and synthesis of genetic evidence have traditionally relied on large-scale, human expert-driven curation efforts that demand substantial time, resources, and specialized domain knowledge^8,9^. Recent advances in large language models (LLMs) provide an opportunity to fundamentally reshape how heterogeneous genetic and functional genomics evidence is collected and deployed in drug development pipelines^10–12^. In principle, LLMs can rapidly integrate information across diverse biomedical resources, reason over heterogeneous lines of evidence and substantially expand the scale, coverage, and accessibility of genetic insights that inform target prioritization and clinical trial decision. Despite this promise, rigorous evaluation is lacking. This gap limits practical pathways for deploying AI-powered genetic evidence in drug development pipelines, as it remains unclear how closely LLM-derived genetic evidence aligns with trusted human expert-curated resources, or whether such models can meaningfully inform clinical success.

To address these important questions, we develop a novel domain-grounded instruction framework to systematically evaluate the performance of GPT-5 in generating genetic evidence for drug development, and we benchmark its outputs against a recent, comprehensive human expert-curated study that reflects years of effort and has become a standard resource in the field^1^. Using 13,022 drug target-indication (T-I) pairs that reached at least Phase I of clinical development since 2000 as labeled data, we evaluate whether LLM-derived genetic evidence is associated with a higher probability of clinical success. To the best of our knowledge, this constitutes the most comprehensive benchmark and analysis of its kind. We show that GPT-5 achieves reliable and methodologically sound performance for automated genetic evidence generation, and provides important advantages in coverage, scalability, and interpretability relative to traditional human expert curation-based approaches.

Building on these findings, we introduce TigerAI (Target-Indication Genetic Evidence Repository through Artificial Intelligence; https://tigerai.bio/), one of the first platforms for AI-powered genetic evidence with two complementary functions (**Fig. 1A**). First, recognizing the rapid evolution of state-of-the-art LLMs, TigerAI provides a rigorous benchmarking framework for systematic comparison of LLMs using large-scale drug development data resources. This framework enables quantitative assessment of model performance and reliability over time. Second, TigerAI provides an easy-to-use interface that delivers AI-powered genetic evidence to researchers and drug developers, produced with rigorously evaluated models and domain-grounded instructions. In summary, our findings and the TigerAI infrastructure support the emerging role of LLMs in drug target discovery, establish a practical pathway for evaluating AI-generated genetic evidence at scale, and highlight the potential of AI to advance understanding of the genetic determinants of clinical success.

**Fig. 1.**
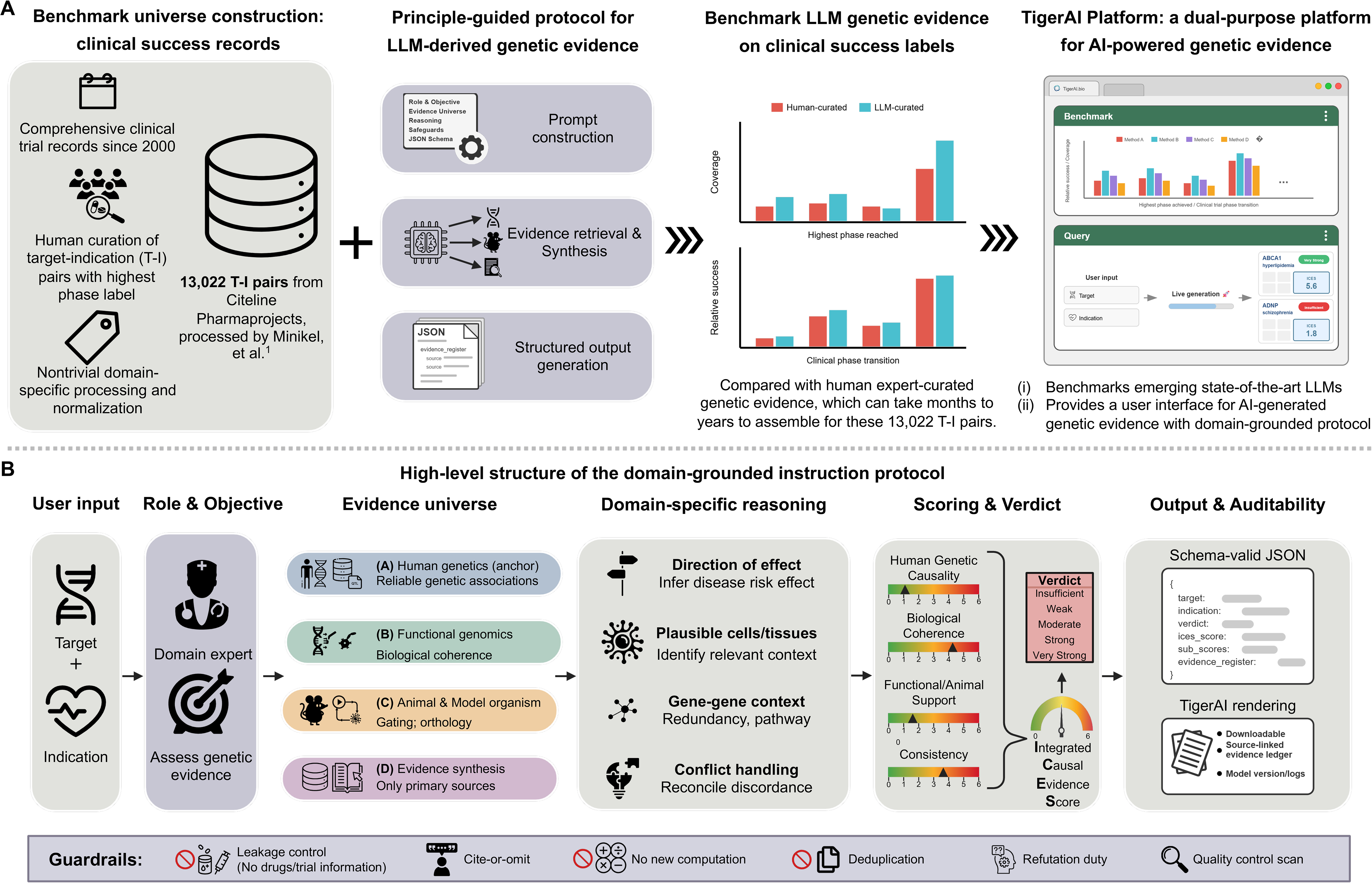
Study overview and domain principle-guided instruction framework. (**A**) Schematic of the study design and TigerAI, a dual-purpose platform for AI-powered genetic evidence. TigerAI (i) benchmarks emerging state-of-the-art LLMs on a large drug development target-indication (T-I) resource and (ii) provides a user-accessible interface for querying structured, source-traceable AI-generated genetic evidence for user-specified T-I pairs using our domain-grounded instruction protocol. (**B**) High-level structure of the domain-grounded instruction protocol used to generate AI-powered genetic evidence for each T-I pair. The protocol instantiates a standardized T-I template and guides evidence synthesis using explicit domain principles, including a predefined evidence universe, domain-specific reasoning requirements, an evidence ledger linking claims to indexed sources, a qualitative-only scoring rubric, model quality safeguards, and a strict machine-readable JSON output organized by evidence sub-category.

### Domain-grounded instructions for AI generation of genetic evidence

We introduced a novel domain-grounded instruction framework (**Methods**) that standardizes LLM-derived genetic evidence to meet domain principles (**Fig. 1B**). For every query, we first instantiate a T-I pair (that is, a specific gene target and a specific indication), ensuring that all subsequent synthesis is conditioned on a clearly defined biological question. We then state in plain language what the model is expected to do: act as a domain expert in human genetics and functional genomics for the relevant therapeutic area and determine whether the gene target is likely to have a causal, human-relevant role in the disease indication. We also explicitly prohibit the use of drug-, therapy-, or clinical trial-related information, preventing potential negative impact from downstream drug development outcomes on LLM reasoning and keeping evidence grounded in upstream genetics and genomics.

To guide genetic evidence gathering, we defined an evidence universe and required the LLM to collect and summarize evidence across four channels that reflect established genetic target validation practice and curated resources. These channels include (i) human genetic evidence as the primary anchor (for example, genome-wide association studies, fine-mapped association signals, variant-to-gene links, molecular quantitative trait locus [QTL] colocalization, rare-variant burden, and phenome-wide association studies); (ii) functional genomics evidence to establish biological coherence (such as tissue and cell-type expression, regulatory/epigenomic context, perturbation evidence, pathways, and interaction networks); (iii) animal and model organism evidence (such as knockout or knock-in phenotypes, under conservative orthology quality requirement, and phenotype alignment criteria); and (iv) evidence synthesis for breadth that can be clearly labeled and verified against the above primary resources. Deduplication and resource-quality preferences are emphasized to reduce inflated support from mirrored databases and to prioritize curated and peer-reviewed resources.

After evidence assembly, we require the LLM to apply domain-specific reasoning principles that mirror how domain experts evaluate genetic causality^13–16^. In general, these principles encompass assessment of the direction of genetic effect; identification of relevant cell types or tissues of action; consideration of gene-gene interactions and network context; interpretation of functional and animal evidence; and principled handling of conflicting findings when previous studies disagree. To maintain causal interpretability, the instructions enforce strict leakage control to exclude drug-related information and emphasize a genetic causality first framework, in which strong conclusions are drawn only when multiple orthogonal lines of evidence converge.

Concretely, the LLM first infers the direction of genetic effect, whether increased or decreased target activity is associated with higher or lower disease risk or severity, prioritizing convergent signals from human genetic studies. It then identifies plausible cell types or tissues in which the association is biologically interpretable, integrating expression and regulatory context to assess cellular plausibility. To reduce misassignment at complex loci, the model draws on its knowledge to evaluate broader biological context, including paralogs, functional redundancy, pathway membership, and network neighbors. The LLM is further required to explicitly consider and reconcile conflicting evidence, such as alternative candidate genes at associated loci, tissue-specific discordance or assay-related artifacts, rather than smoothing over inconsistencies. Finally, cross-ancestry heterogeneity will be documented and contextualized but not treated as contradictory unless there is explicit refutation, aligning the synthesis with best practices for interpreting human genetic causality.

To ensure transparency and auditability, we enforce an evidence-ledger principle: every substantive claim made by the LLM must be supported by at least one indexed source entry, that is, linked to a primary research study or a curated database record. We further standardize interpretation through a qualitative-only scoring framework, in which evidence is strictly summarized across four interpretable categories by fully leveraging LLM’s internal reasoning ability: (i) human genetic causality, (ii) biological coherence, (iii) functional and animal support, and (iv) consistency across evidence types. Then, these components are integrated into an integrated causal evidence score (ICES; range 0-6), which is mapped to a small set of verdict levels (insufficient, weak, moderate, strong, and very strong), using prespecified thresholds to enable consistent calibration across targets and indications. Unless otherwise noted, we define genetic support as ICES ≥ 4.0, corresponding to strong or very strong verdicts.

In addition, we incorporate explicit LLM reliability safeguards and a strict output contract to support benchmarking at scale. Reliability safeguards include a “cite or omit” rule that requires the LLM to corroborate major causal and direction-of-effect statements, and an explicit duty to search for counterevidence and constraints that prevent *de novo* computation. The LLM is also instructed to flag potential technical pitfalls (for example, protein QTL assay artifacts and platform discordance) when relevant. All outputs are returned in a strict, machine-readable JSON structure, organized by evidence category, enabling systematic comparison with curated resources and consistent evaluation as LLMs evolve.

### Benchmarking AI-generated genetic evidence against expert curation

We applied the domain-grounded instruction framework described above to GPT-5 and generated AI-powered genetic evidence for 13,022 T-I pairs from drug development records that reached at least Phase I. These pairs were drawn from a previously curated dataset that filtered Citeline Pharmaprojects for monotherapy programmes added since 2000 with an annotated highest phase reached (Phase I-III or Launched). Each programme was mapped to a human gene target (typically the gene encoding the drug target protein) and an indication defined using the Medical Subject Headings (MeSH) ontology. Most analyses reported below focus on comparing GPT-5 outputs with a recent exhaustive human expert-curated analysis of the same drug development resource^1^, which integrated evidence from multiple major genetics and genomics databases, and reported that drug mechanisms with genetic support have a 2.6-fold higher probability of success than those without (**Methods**). We also applied the same instruction framework to additional LLMs and present LLM model comparisons in a later section.

On this comprehensive drug development benchmark of 13,022 T-I pairs, genetic evidence derived by GPT-5 showed associations with clinical success that were broadly comparable to those obtained from human expert-curated resources. Across clinical stages, GPT-5 preserved similar enrichment patterns across the development pipeline and provided comparable or higher coverage of genetically supported T-I pairs than the human expert-curated baseline. In Phase I (*n* = 2,569), genetic support was identified for 93 T-I pairs (3.6%) by human expert curation versus 140 (5.4%) by GPT-5; in Phase II (*n* = 4,153), 153 (3.7%) versus 210 (5.1%); and in Phase III (*n* = 665), 30 (4.5%) versus 28 (4.2%). The largest difference was observed among launched programmes (*n* = 1,519), where GPT-5 identified 312 supported pairs (20.5%), compared with 189 (12.4%) by expert curation (**Fig. 2A**). Consistent with previous study^1^, genetic support defined by either approach was associated with increased relative success (RS) across clinical trial phase transitions, with GPT-5 yielding slightly larger RS than human expert curation: 1.10 versus 1.08 (Phase I to II), 1.68 versus 1.57 (Phase II to III), 1.40 versus 1.28 (Phase III to Launched), and 2.74 versus 2.63 for the overall Phase I to Launched transition (**Fig. 2B**). Here RS was defined as the ratio of the probability of success with identified genetic support to the probability of success without genetic support (**Methods**). These results indicate that GPT-5 derived genetic evidence recovers the well-established link between human genetic support and clinical success, supporting the promise and reliability of GPT-5 for this scientific task.

**Fig. 2.**
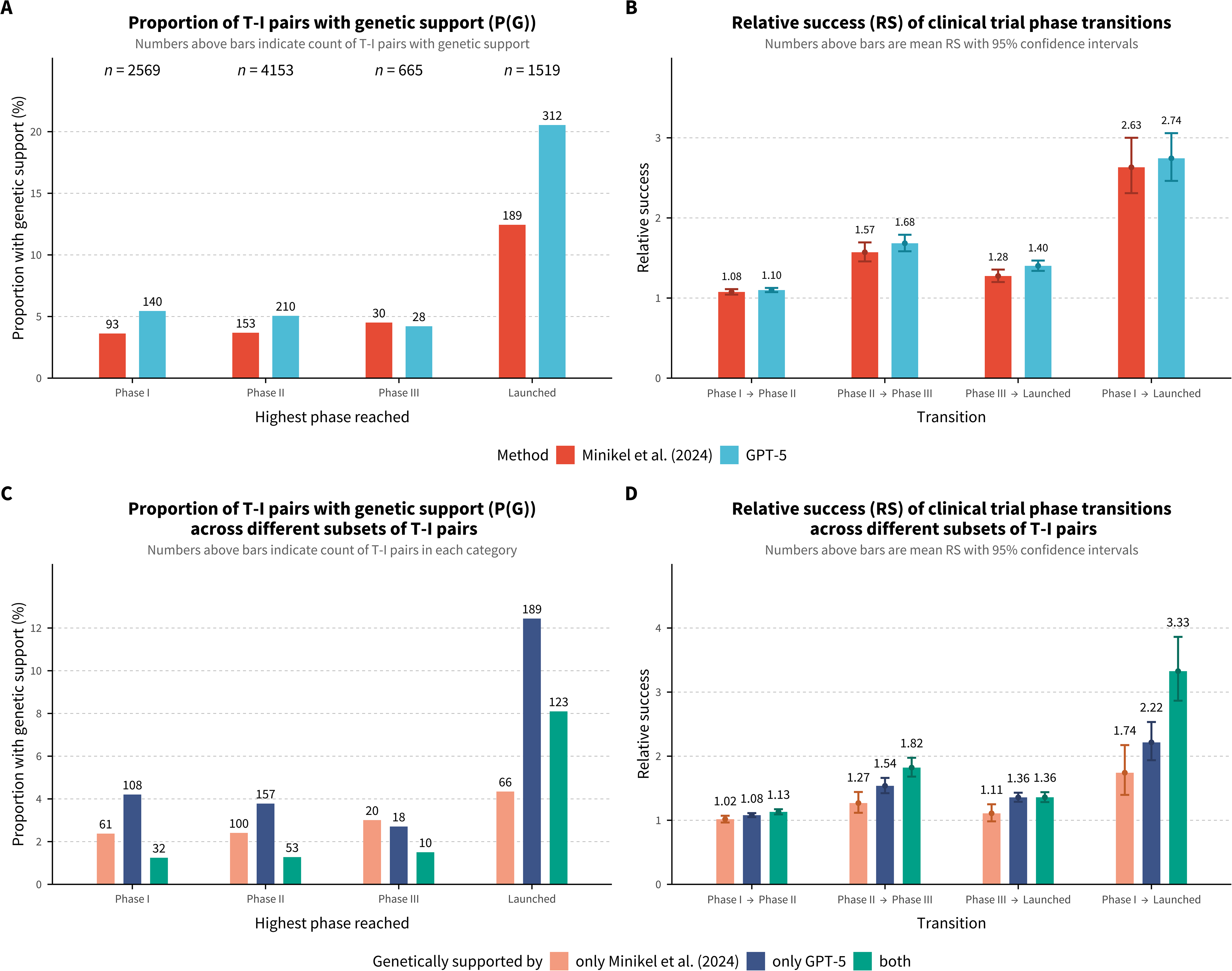
Comparing GPT-5-derived genetic evidence with human expert curation. (**A**) Proportion of target-indication (T-I) pairs classified as genetically supported, P(G), stratified by highest clinical phase reached (Phase I-III and Launched), comparing GPT-5-derived evidence with exhaustive human curation by Minikel, et al. ^1^. Numbers above bars indicate counts of T-I pairs meeting criteria; *n* values indicate total T-I pairs per phase. (**B**) Relative success (RS) of clinical trial phase transitions, defined as the ratio of transition probability for genetically supported T-I pairs to that of unsupported pairs. RS and the associated 95% confidence intervals were calculated using the Katz log method for all transitions, including Phase I to Launched. Error bars represent 95% confidence intervals. (**C**) P(G) stratified by annotation agreement: T-I pairs identified exclusively by human curation, exclusively by GPT-5, or by both methods. Numbers indicate counts per category. (**D**) RS stratified by agreement group (human-only, GPT-5-only, and both) across the same transitions as in (**B**), with the strongest enrichment (RS = 3.33) observed when both methods support the same T-I pair. Analyses use 13,022 T-I pairs from monotherapy programmes added since 2000 that reached at least Phase I; GPT-5 evidence was generated using the domain-grounded instruction described in this study.

Stratifying T-I pairs by agreement between GPT-5 and human curation showed both substantial overlap and complementary genetic evidence. Among launched programmes, 66 T-I pairs were supported only by human curation, 189 only by GPT-5, and 123 by both approaches (**Fig. 2C**). Notably, RS was highest for pairs supported by both approaches, intermediate for GPT-5-only support, and the lowest for human-only support: for overall Phase I to Launched, RS was 1.74 (human-only), 2.22 (GPT-5-only), and 3.33 (both) (**Fig. 2D**). These stratified results suggest that concordant support marks the highest-confidence subset, while additional T-I pairs only supported by GPT-5 remain enriched for clinical success. When we considered T-I pairs supported by either method, coverage increased to 378/1,519 (24.9%) among launched programmes while maintaining an overall RS of 2.64 (**Fig. S1**). Collectively, these results support that GPT-5 provides additional genetic evidence that is not captured in commonly used human expert-curated data resources.

To further analyze differences between GPT-5 and expert curation, we stratified T-I pairs by indication complexity, defined by the number of therapeutic areas to which each MeSH term maps (single versus multiple therapeutic areas). For launched programmes, GPT-5 showed moderately higher genetic support coverage than human curation for indications spanning multiple therapeutic areas (92 vs 102 supported pairs), together with a larger increase in overall Phase I to Launched RS (2.64 vs 3.28). In contrast, for indications assigned to a single therapeutic area, GPT-5 substantially increased coverage (97 vs 210 supported pairs) while the overall Phase I to Launched RS remained similar (2.52 vs 2.53) (**Fig. S2A-S2B**). Decomposing coverage by agreement group further clarified how complexity interacts with overlap between the two approaches: among launched programmes, multi-area indications comprised 28 human-only, 38 GPT-5-only, and 64 jointly supported pairs, whereas single-area indications comprised 36 human-only, 151 GPT-5-only, and 39 jointly supported pairs, indicating that the largest incremental coverage from GPT-5 occurs for single-area indications (**Fig. S2C**). In both groups, RS was highest for jointly supported pairs, intermediate for GPT-5-only support, and lowest for human-only support (**Fig. S2D**). We also contrasted oncology versus non-oncology programmes. GPT-5 identified a higher proportion of genetically supported launched T-I pairs than human expert curation in both groups, with a more pronounced increase in oncology than in non-oncology (**Fig. S3**). Overall, these stratified analyses suggest that compared to human expert-curated genetic support, GPT-5 expands genetic support coverage across indication complexity and therapeutic context strata to varying degrees, while generally preserving, and in some settings strengthening, the enrichment for clinical success.

### Dissecting AI-generated genetic evidence by therapeutic areas

We next evaluated GPT-5 performance within individual therapeutic areas. Across categories, the pattern of genetic support coverage derived from GPT-5 was broadly consistent with that observed under human expert curation (**Fig. 3A**): across clinical stages, GPT-5 generally identified comparable coverage profiles by category. When restricted to launched programmes, the proportion of genetically supported T-I pairs identified by GPT-5 and by human curation was positively correlated across therapeutic areas (Correlation *ρ* = 0.62, **Fig. 3B**), indicating broad concordance. Differences were more pronounced in a few categories, including endocrine and oncology, where GPT-5 identified higher coverage, and psychiatry, where expert curation identified higher coverage. In contrast, several categories, such as metabolic, congenital, and hematology, showed near-identical coverage level between GPT-5 and expert curation. These categories also showed higher-than-average overlap in genetic support between human expert curation and GPT-5, and GPT-5 consistently contributed a large proportion of novel, genetically supported T-I pairs across therapeutic areas (**Fig. S4**)

**Fig. 3.**
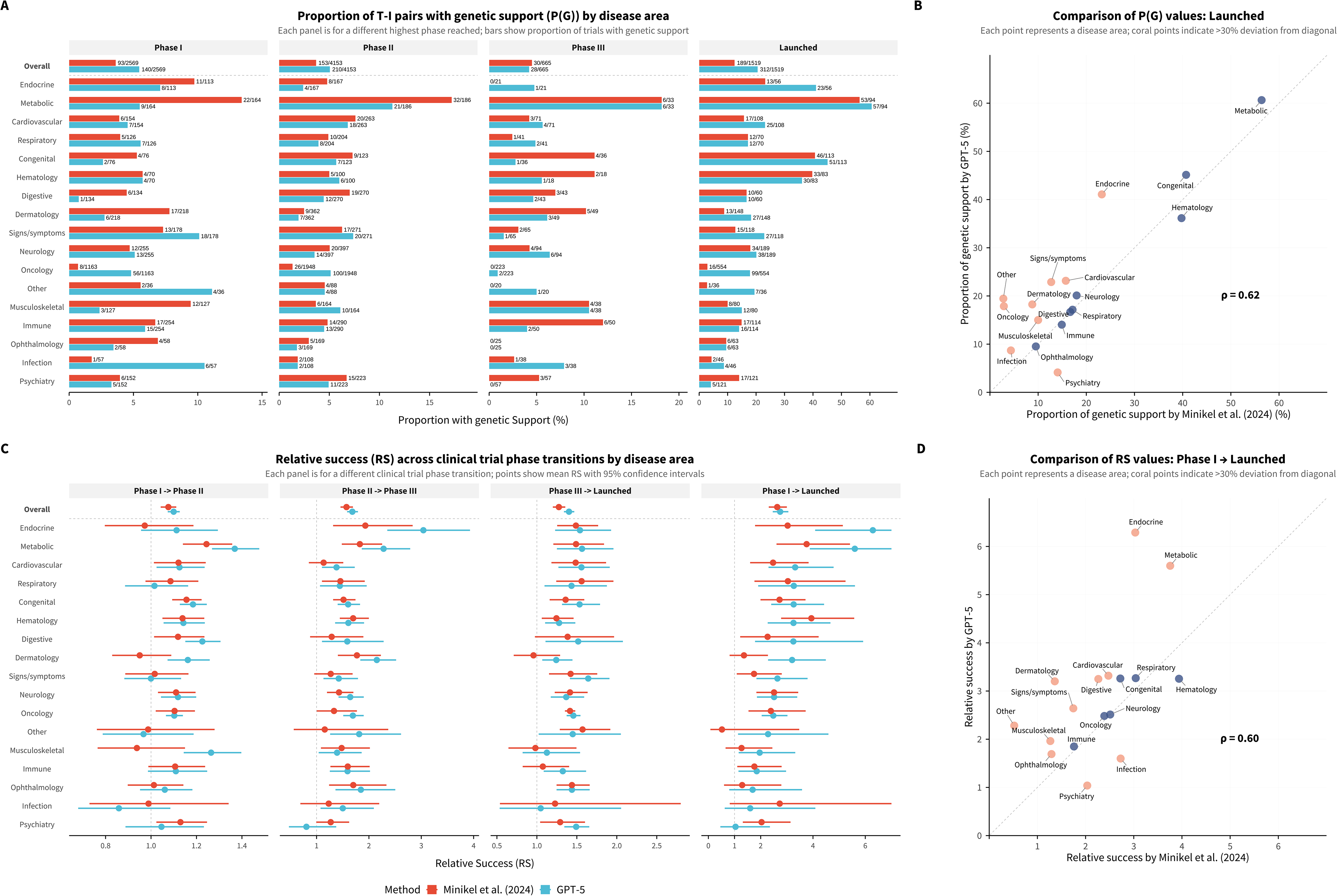
Therapeutic area-specific patterns of GPT-5-derived genetic evidence. Comparison of genetic support metrics between human expert curation by Minikel, et al. ^1^ and GPT-5 across therapeutic areas. (**A**) Proportion of target-indication (T-I) pairs with genetic support, P(G), by therapeutic area and the highest clinical phase reached. Numbers adjacent to bars indicate counts of genetically supported T-I pairs relative to total pairs. (**B**) Scatterplot comparing P(G) values at the Launched phase between GPT-5 and human curation across 17 therapeutic areas. Coral points indicate therapeutic areas with >30% deviation from the diagonal (y=x). Spearman correlation coefficient (ρ) is shown. (**C**) Relative success (RS) of clinical trial phase transitions by therapeutic area, comparing GPT-5 (blue) and human expert curation (red) for assigning genetic support. RS is defined as the ratio of transition probability for genetically supported T-I pairs to that of unsupported pairs. Each panel represents a different phase transition: Phase I to Phase II, Phase II to Phase III, Phase III to Launched, and Phase I to Launched. Points indicate mean RS values; error bars represent 95% confidence intervals calculated using the Katz log method. Therapeutic areas are ordered by descending RS for the Phase I to Launched transition. (**D**) Scatterplot comparing RS values for the Phase I to Launched transition between GPT-5 and human curation. Coral points indicate therapeutic areas with >30% deviation from the diagonal. Despite methodological differences, both approaches show moderate-to-strong correlation across therapeutic areas (ρ=0.62 for P(G) and ρ=0.60 for RS), with the largest discrepancies observed in metabolic, endocrine, and congenital therapeutic areas.

We then assessed category-specific differences in the RS associated with genetic support. For most therapeutic areas, GPT-5 derived genetic evidence yielded RS estimates broadly comparable to those obtained from human-curated support across Phase I to II, Phase II to III, Phase III to Launched, and the overall Phase I to Launched transitions (**Fig. 3C**). Focusing on the aggregate Phase I to Launched transition, RS estimates derived from GPT-5 and from expert curation were positively correlated across therapeutic areas (Correlation *ρ* = 0.60, **Fig. 3D**), indicating that categories showing stronger enrichment under human expert curation tended to show similar enrichment under GPT-5. Differences were more apparent in several areas, including endocrine and metabolic diseases, where GPT-5 tended to yield higher RS estimates, whereas expert curation produced higher RS in psychiatry and infection. Several categories lay close to the identity line, indicating close agreement between GPT-5 and human expert curation, including hematology, respiratory, congenital, neurology, oncology, and immune. In summary, these disease-stratified analyses suggest that GPT-5 generated genetic evidence largely preserves disease therapeutic area-specific patterns observed in human expert-curated resources. Among launched programmes, GPT-5 often provides comparable or higher coverage and similar or higher RS across most therapeutic domains.

### Evidence-dimension interpretability and ICES thresholding

After benchmarking against human expert curation, we next focused on the GPT-5 generated outputs to understand how genetic support is distributed across the scoring system dimensions, beyond the single ICES. This analysis provides insights into the composition and interpretability of the evidence produced by our domain-grounded instructions. We examined the four interpretable evidence categories: human genetic causality, biological coherence, functional and animal support, and consistency across evidence types. Across all four categories, we observed a consistent phase-dependent enrichment of genetically supported T-I pairs, but with distinct coverage profiles (**Fig. 4A**). Biological coherence was the most inclusive category at every stage (for example, 364/2,569 (14.2%) in Phase I, 545/4,153 (13.1%) in Phase II, 82/665 (12.3%) in Phase III, and 529/1,519 (34.8%) among launched programmes), whereas human genetic causality was the most selective (112/2,569 (4.4%), 162/4,153 (3.9%), 24/665 (3.6%), and 244/1,519 (16.1%), respectively). Functional and animal support, as well as evidence consistency, showed intermediate coverage with similar phase-dependent patterns.

**Fig. 4.**
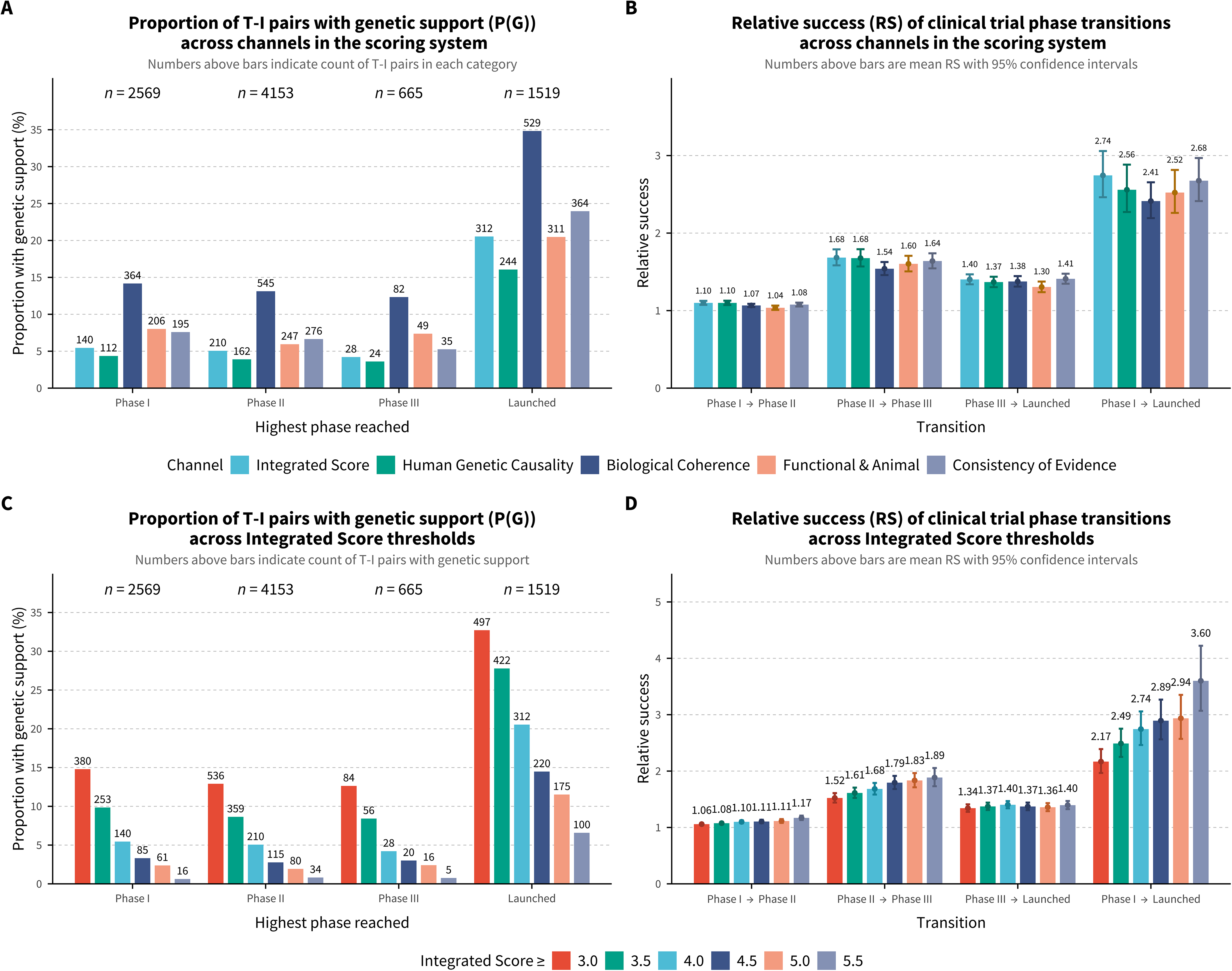
Evidence channels and integrated causal evidence score thresholding for GPT-5-derived genetic evidence. (**A**) Proportion of target-indication (T-I) pairs with genetic support, P(G), across four channels of the scoring system, stratified by highest clinical phase reached. Genetic support is defined as a channel score ≥ 4.0. Numbers above bars indicate counts of T-I pairs meeting criteria; *n* values indicate total T-I pairs per phase. (**B**) Relative success (RS) of clinical trial phase transitions, defined as the ratio of transition probability for genetically supported T-I pairs to that of unsupported pairs. RS and 95% confidence intervals were calculated using the Katz log method for all transitions, including Phase I to Launched. Error bars represent 95% confidence intervals. (**C**) Proportion of T-I pairs with genetic support at varying integrated causal evidence score (Integrated Score) thresholds (≥3.0 to ≥ 5.5), stratified by highest clinical phase reached. (**D**) RS for each subset defined in (**C**). As the Integrated Score threshold becomes more stringent, fewer T-I pairs qualify as having genetic support, but the resulting subsets exhibit progressively higher RS values, consistent with an expected trade-off between coverage and strength of genetic support. Error bars represent 95% confidence intervals.

Across phase transitions, genetic support defined by each evidence category was associated with increased RS, with broadly similar qualitative patterns and expected differences in magnitude by transition (**Fig. 4B**). Aggregating across the full pipeline (Phase I to Launched), RS estimates across the four evidence categories ranged from 2.41 to 2.68. Biological coherence, despite providing the broadest genetic coverage, showed the lowest RS among the four channels. The integrated score (that is, the ICES) yielded the largest enrichment (RS = 2.74). These results indicate that, as a composite summary measure, the ICES used in our primary analyses reflected a nuanced and reasoned assessment that balanced evidence across the four information channels, yielding the strongest overall RS signal.

Furthermore, the ICES (range 0-6) is designed as a tunable summary of evidence strength, with higher values intended to reflect stronger and more convergent genetic support. To assess its practical flexibility, we evaluated how the fraction of genetically supported T-I pairs and their associated RS vary across ICES thresholds. As expected, stricter thresholds produced a monotonic reduction in coverage across all clinical stages (**Fig. 4C**). Among launched programmes (*n* = 1,519), an ICES threshold of 3 yielded 497 supported T-I pairs while maintaining meaningful enrichment (RS = 2.17) (**Fig. 4D**), consistent with a moderate threshold capturing broader yet still informative genetic support. Coverage then decreased monotonically with stricter ICES thresholds, to 312 at ICES ≥ 4.0 (threshold used in the primary analyses), and 175 at ICES ≥ 5.0. In parallel, for the Phase I to Launched transition, RS increased with stricter thresholds, rising to 2.74 at ICES ≥ 4.0 and 2.94 at ICES ≥ 5.0. These results suggest that ICES captures the strength of genetic evidence and highlight a practical trade-off between breadth and stringency: higher thresholds yield smaller sets of supported T-I pairs with stronger RS, whereas lower thresholds provide broader coverage with more moderate RS.

### Operationalizing AI-generated genetic evidence with the TigerAI platform

After illustrating that GPT-5 yields reliable genetic evidence for clinical success, we were motivated to build a public-facing platform to translate this capability into a practical resource for drug target assessment. The resulting web platform, TigerAI (https://tigerai.bio/), is novel in two complementary aspects: it enables standardized evaluation of emerging state-of-the-art LLMs on a large drug development benchmark dataset, and it provides on-demand, real-time genetic evidence generated by validated LLMs under our domain-grounded instructions. Unlike traditional human-curated databases, which are inherently static and often cover genetic information only for a limited subset of T-I pairs in drug development, TigerAI can generate structured, query-specific evidence summaries for user-specified T-I pairs, while preserving traceability to underlying sources and providing a transparent breakdown across evidence dimensions. Real-time query-specific generation is also a pragmatic design choice in the AI setting. Because LLM capabilities and versions evolve rapidly, precomputed LLM-derived databases may risk becoming outdated quickly, and maintaining multiple versions across models would be costly. For example, running our domain-grounded instruction incurs a nontrivial per-query cost (approximately $0.12; **Table S1**). Precomputing outputs exhaustively across thousands of indications and about 20,000 protein-coding genes would therefore be inefficient and difficult to sustain. By generating results at query time, TigerAI provides a scalable pathway for using AI-generated genetic evidence in practice.

On the LLM benchmarking side, TigerAI leverages one of the largest curated datasets from the drug development pipeline to enable efficient and standardized evaluation of model performance. New models or LLM-based methods, tested either by us or by model developers, can be incorporated by running them through the same domain-grounded instruction protocol, allowing their outputs to be analyzed in parallel with human expert curation, as illustrated for GPT-5 in **Figures 2-4**. Furthermore, by holding the benchmarking data, prompts, and scoring framework fixed, TigerAI enables more fair comparisons across LLMs, making it possible to interpret their differences in genetic coverage and RS. For illustration purpose, we compared GPT-5 with GPT-4o. Across clinical phase stages, both models showed the expected stage-dependent increase in genetically supported T-I pairs, but with systematic differences in coverage (**Fig. 5A**). GPT-4o identified substantially more supported T-I pairs than GPT-5 at Phase I-III (over two-fold), but this difference attenuated for launched programmes, where the increase was smaller and fell below two-fold. Despite these coverage differences, RS estimates across phase transitions were broadly similar (Phase I to Launched RS 2.74 for GPT-5 versus 2.60 for GPT-4o), although GPT-4o’s RS was lower than that obtained from human curation (**Fig. 5B**). At the output level, GPT-5 and GPT-4o showed moderate concordance in qualitative verdicts and ICES values. Both models agreed on many T-I pairs classified as strong or very strong, but GPT-4o tended to assign higher scores more broadly, whereas GPT-5 exhibited a more conservative distribution. Notably, a substantial subset of T-I pairs labeled strong by GPT-4o were assigned moderate, weak, or insufficient evidence by GPT-5, contributing to the coverage gap between models (**Fig. 5C**). Consistent with this pattern, GPT-4o scores were generally higher than GPT-5 across ICES and the individual evidence channels, with positive correlations between the models (**Figs. 5D-5H**). This pattern suggests differences in how GPT-4o and GPT-5 interpret and calibrate target-indication evidence, with GPT-5 applying a more conservative evaluation that more closely aligns with the human expert-curated baseline in this task. We will continue to expand LLM benchmarking on TigerAI by incorporating more recent models (such as Gemini 3) and providing more comprehensive perspectives for model comparison.

**Fig. 5.**
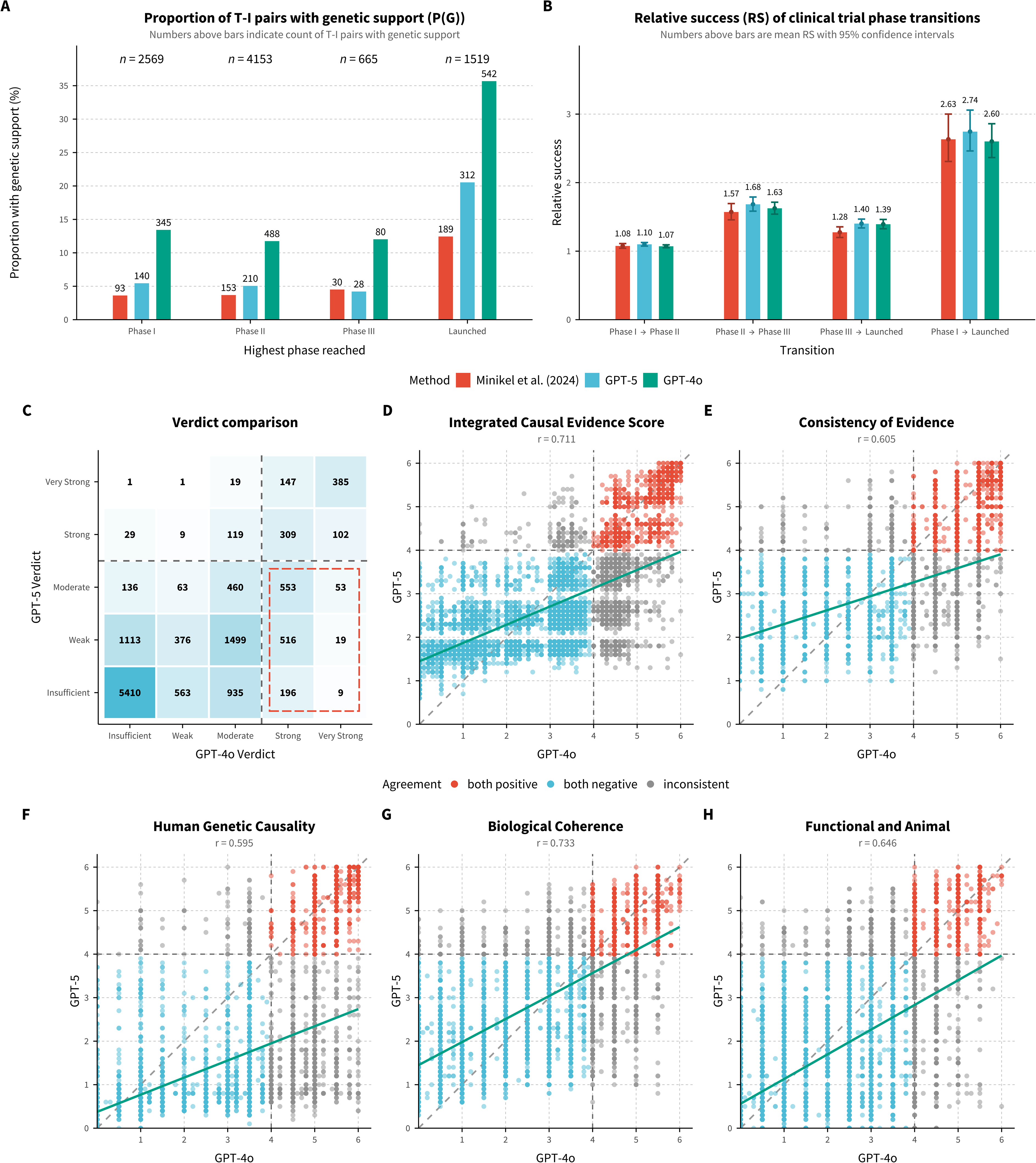
Cross-model benchmarking of AI-generated genetic evidence: GPT-5 versus GPT-4o. (**A**) Proportion of target-indication (T-I) pairs with genetic support, P(G), stratified by highest clinical phase reached and classification method. T-I pairs were classified using human expert curation by Minikel, et al. ^1^, GPT-5, or GPT-4o. Numbers above bars indicate counts of T-I pairs with genetic support. (**B**) Relative success (RS) of clinical trial phase transitions, stratified by classification method. Numbers above bars indicate mean RS values; error bars represent 95% confidence intervals. (**C**) Confusion matrix comparing verdict labels assigned by GPT-4o and GPT-5. Cell values denote counts of T-I pairs in each verdict combination. The asymmetry between counts below the diagonal (4,445) versus above (1,637) indicates that GPT-4o tends to assign higher-evidence verdicts than GPT-5. (**D-H**) Agreement of score components between GPT-4o and GPT-5, shown as scatterplots for the integrated causal evidence score (**D**) and the four evidence dimensions: consistency (**E**), human genetic causality (**F**), biological coherence (**G**), and functional/animal support (**H**). Pearson correlation coefficients (*r*) are reported. Points are colored by verdict agreement (both supported: strong/very strong; both unsupported: insufficient/weak/moderate; or discordant), dashed diagonals indicate perfect agreement, and solid lines show linear regression fits. GPT-4o scores show visible discretization (clustering at 0.5 increments), consistent with coarser score calibration relative to GPT-5.

On the user-facing side, TigerAI provides a user-friendly interactive interface for researchers to evaluate genetic evidence for a specific T-I pair using our domain-grounded instruction protocol. Each query returns a standardized, downloadable report that summarizes the overall strength of AI-generated genetic support and decomposes it into interpretable dimensions, accompanied by a source-linked evidence register that enables users to review supporting details, such as the primary studies and curated records underlying the assessment. Users can also compare outputs across multiple benchmarked LLMs (for example, GPT-5 and GPT-4o) and model options (such as reasoning level). Collectively, TigerAI enables standardized head-to-head benchmarking, exposing differences in calibration and evidence stringency across LLMs under identical evaluation conditions. In parallel, it provides transparent, real-time access to AI-generated genetic evidence, providing a practical alternative to static, labor-intensive genetic evidence curation.

## Discussion

In this study, we assessed whether modern LLMs can generate genetic evidence that is informative for clinical success in drug development. We developed a domain-grounded instruction framework that uses structured prompting to standardize evidence collection, reasoning, and reporting. Using a large benchmark of T-I pairs from Citeline Pharmaprojects, we show that GPT-5 generated genetic evidence is broadly comparable to human expert curation in its ability to inform higher probability of clinical success, while offering substantially broader coverage than traditional curated resources. We further examined key properties of the GPT-5 outputs, including how performance varies with indication complexity and therapeutic area, the contribution of interpretable evidence dimensions, and the trade-off between coverage and RS under different evidence thresholds. To the best of our knowledge, this drug development collection represents the most comprehensive benchmark to date, and our evaluation is the first to demonstrate that LLM-derived genetic evidence can provide reliable signals for clinical success. Building on these analysis findings, we introduced TigerAI, a public-facing platform that systematically benchmarks emerging LLMs and delivers real-time, query-specific genetic evidence generated under rigorously domain-grounded protocols, supporting the responsible use of AI-enabled genetic evidence in the community.

A first key contribution of this work is the development of a principled instruction framework for generating and evaluating AI-derived genetic evidence. We anchor evidence generation in established domain practice^13–16^, require traceability to underlying resources, and assess performance on a clinically meaningful benchmark dataset. Under these constraints, AI-generated genetic evidence recovers key signals previously identified through extensive manual curation^1^ while identifying additional T-I pairs that remain enriched for success. Our findings suggest that, when carefully specified and systematically evaluated, current LLM-based AI systems can meaningfully support genetic evidence synthesis.

Beyond these empirical analyses and findings, a second major contribution is the development of TigerAI, an AI-powered platform that supports systematic and transparent progress in this area. Because LLM capabilities evolve rapidly, one-off evaluations can be quickly outdated. TigerAI addresses this by enabling newly released models to be benchmarked under the same fixed protocol as they emerge, allowing direct, comparable assessments over time. This infrastructure moves the field beyond isolated comparisons toward ongoing, reproducible evaluation of AI systems for genetic evidence generation. TigerAI also introduces a real-time evidence paradigm that complements traditional curated resources, which are static and often updated on slower cycles. Using our domain-expert instruction protocol, researchers can query a specific T-I question and obtain a structured, source-linked evidence summary on demand. Importantly, TigerAI contextualizes these outputs with benchmarking results derived from the drug development pipeline, giving users an explicit performance reference when interpreting the strength of LLM-generated evidence.

As an early large-scale study examining the potential of LLMs to synthesize genetic evidence for drug development, our work has several limitations and highlights key open questions for future studies aimed at building next-generation AI systems for genetic evidence synthesis. First, despite imposing strict domain-specific constraints on admissible evidence and requiring resource-linked support, the quality of model outputs remains highly dependent on the completeness of the underlying literature and resources used for LLM training, and may therefore inherit their biases and gaps. Our analysis suggests that GPT-5 and GPT-4o are already reliable for this task, with GPT-5 more closely aligning with the human expert-curated baseline. By comparison, GPT-3.5 did not produce meaningful results with our domain-grounded instructions (**Methods**). Based on these observations, we expect continued performance gains from future state-of-the-art LLMs.

Moreover, although our protocol explicitly excludes drug- and clinical trial-related information, complete absence of leakage in model behavior cannot be guaranteed in theory. To mitigate this concern, we systematically scanned the LLM outputs across all 13,022 T-I pairs using a curated dictionary of terms commonly associated with drug development and clinical trial contexts. Only 2% (253) of T-I pairs were flagged as containing potentially suspicious evidence suggesting possible mixing with drug- and trial-related terminology. Upon detailed manual review of all 253 pairs, we found no evidence of information leakage; instead, these cases reflected legitimate biological terminology unrelated to therapeutic contexts, or explicit model self-reflection describing the deliberate exclusion of therapy- and response-related literature to avoid confounding. These results substantially reduce concerns about potential data leakage and suggest that modern LLMs can reliably follow our domain-grounded instructions to generate genetic evidence without incorporating non-genetic contexts, such as unwanted drug or trial-related information.

In addition, owing to the generative nature of LLMs, model outputs may contain hallucinated content and exhibit variability across prompting configurations. To assess robustness, we conducted additional analyses across all 13,022 T-I pairs and found that GPT-5 was generally stable for this task, showing overall high consistency when rerun at scale using different APIs (**Fig. S5**). To support auditing and transparency, each TigerAI user report will explicitly label the model version, preserve prompt settings, and provide structured links to the underlying literature supporting the generated genetic evidence. Finally, the development of TigerAI and other AI-powered platforms does not obviate the need for human expertise or established human curated resources^1,8,9,17^: AI-generated genetic evidence should be viewed as decision support that complements, rather than replaces, expert judgment and ongoing data integration efforts in genetic target assessment, and the two approaches can be mutually reinforcing.

Looking ahead, several directions could further expand the scope and impact of AI-based genetic analysis in drug discovery and beyond. Continued advances in LLM capability, such as stronger and more efficient reasoning^18,19^, prompt optimization^20–22^, and more reliable benchmarking in high-quality biomedical data resources, are expected to improve both the accuracy and interpretability of evidence synthesis. Beyond text-based knowledge, multimodal data resources and foundation models that jointly integrate genetic variation^8,9^, molecular phenotypes^23^, imaging^24^, and biological networks^25^ may better link genetic association signals to underlying mechanisms. In parallel, agentic AI systems^26–28^ that iteratively retrieve evidence, improve prompt, propose follow-up analyses, and refresh conclusions as new data emerge may provide a path toward more adaptive, continuously updated genetic evidence generation. Such AI systems could support multiple genetic applications in drug development^29^, including side effect risk prediction^30,31^, drug target prioritization^32^, and insights into clinical trial termination^33^. From a translational perspective, these AI advances could also meaningfully shape how genetic evidence is used in drug repurposing^34,35^, precision therapeutics^36^, and clinical genetics^37–39^. As models, data resources, and evaluation standards mature, integrating these tools into translational workflows in a principled and transparent manner will be essential to realizing their potential to improve clinical success.

## ACKNOWLEDGEMENTS

We are grateful to Matthew R. Nelson, Eric Vallabh Minikel, and Jeffery L. Painter for helpful discussions on the drug development pipeline data used in this study and for insights into their work on data collection and processing. Research reported in this work was supported by National Institute of Mental Health under Award Number R01MH136055; National Institute on Aging under Award Numbers RF1AG082938 and R01AG085581; and National Cancer Institute under Award Numbers R01CA263494, U01CA293883, and P30CA016672; and The U Foundation U-Pilot Award. The content is solely the responsibility of the authors and does not necessarily represent the official views of the National Institutes of Health. We would like to thank Wharton Computing’s Research IT Team for providing computational resources and support that have contributed to these research results.

## Methods

### Domain-grounded instruction and prompt engineering

We formulated genetics-based evidence extraction for each target-indication (T-I) pair as a constrained large language model (LLM) inference task using a fixed developer prompt and a minimal per-pair user prompt (gene and indication). On one hand, the developer prompt explicitly defines the model as an expert of human genetics, restricts admissible evidence to non-interventional human genetics with aligned functional/model support, and explicitly prohibits any use of drug-, therapy-, or clinical trial-related information to mitigate any potential outcome leakage when later benchmarking genetic evidence with clinical development information. On the other hand, the per-query user message supplies only the minimal variables needed for extraction (gene and indication). This separation exploits the API role hierarchy, in which developer messages are prioritized ahead of user messages, to minimize behavioral drift and keep the rubric invariant across T-I pairs. Outputs are returned as a single schema-valid JSON record via Structured Outputs (JSON Schema), enabling scalable downstream analysis of an evidence register, qualitative sub-scores across four channels on a 0-6 scale: (i) human genetic causality, (ii) biological coherence, (iii) functional and animal support, and (iv) consistency across evidence types. We instruct the LLM to qualitatively synthesize sub-scores across the four evidence channels, leveraging its reasoning to form integrated causal evidence score (ICES), which is then mapped to an ordinal verdict. Below, we describe the design of our domain-grounded instruction protocol around four key considerations.

First, we aim to make the assessment scientifically interpretable while remaining auditable. The developer message therefore defines an admissible evidence universe and an explicit synthesis logic. Specifically, the prompt instructs the model to ground its assessment in convergent human genetic anchors (for example, replicated association signals and genetic regulatory evidence that strengthens locus-to-gene assignment and supports a direction of effect), then to evaluate biological coherence using functional genomics and pathway/network context, and finally to incorporate functional and model-organism studies only when they are compatible with human evidence. This design aligns with common practice in locus-to-gene and locus-to-mechanism interpretation, where curated resources such as the GWAS Catalog^40^, Open Targets^8^, eQTL Catalogue^41^, and curated pathway resources (such as Reactome^42^) are used to triangulate causal gene assignment and biological context. The prompt also requires explicit conflict handling: It must represent uncertainty, enumerate credible alternative locus-to-gene assignments and discordant directions of effect, and consider null or refuting evidence rather than forcing a single mechanistic narrative.

Second, we enforce leakage control explicitly in the developer prompt. The prompt prohibits the model from consulting, using, or citing drug- or clinical trial-related information, including drug names, trial outcomes, approvals, therapeutic context, registries and regulatory labels, and instructs it to base its reasoning exclusively on non-interventional human genetics and aligned biological evidence.

Third, the prompt operationalizes the scientific reasoning steps typically required to translate genetic associations into a target-level causal hypothesis. It requires an explicit direction-of-effect statement when support is sufficient, nominates plausibly relevant cells/tissues of action, and incorporates gene-gene interaction context (such as redundancy/paralogy and network/pathway placement) to assess biological plausibility. Functional and animal/model-organism evidence is integrated via a gating rule: such evidence can increase confidence only when orthology is credible and phenotypes align at the phenotype-class level using cross-species phenotype frameworks; otherwise, it is explicitly labeled non-informative rather than treated as implicit support. To reduce genetic evidence inflation from syndication of the same primary result across multiple aggregators, the developer prompt includes an explicit deduplication principle (that is, count each unique primary study/accession once, with mirrored databases noted).

Fourth, we encoded a qualitative, calibration-oriented scoring system that separates evidence dimensions while avoiding false numerical precision. The model returns four qualitative sub-scores on a 0-6 scale (one decimal place): (i) human genetic causality (strength/replication and clarity of causal gene and direction of effect), (ii) biological coherence (tissue/cell relevance and mechanistic context), (iii) functional and animal alignment (rigor and compatibility with human evidence), and (iv) consistency across evidence types (agreement across different evidence channels versus replicated contradictions). The model also returns an ICES (0-6) with a short integration rationale, and the final ordinal verdict is determined solely from pre-specified ICES thresholds (for example, ≥ 4.0 indicating strong and very strong genetic evidence as used in our main analyses) rather than by averaging sub-scores, which makes the decision rule explicit and prevents implicit weighting drift.

Finally, we implemented prompt-level reliability controls and output engineering to improve auditability and large-scale usability. Because LLMs can produce fluent but unsupported content and may fabricate scholarly references, the prompt requires “cite-or-omit,” mandates that major claims (including the main causal claim and direction of effect) be supported by multiple independent sources when available, and includes a “refutation duty” that explicitly prompts the model to consider counterevidence. The prompt also prohibits *de novo* statistical computation and instructs the model to report statistics only as stated in sources, and it includes explicit assay-caveat handling for affinity-based proteomics (such as epitope effects and cross-platform discordance), reflecting known interpretability limitations of protein quantitative trait locus measured by antibody/aptamer platforms. For machine-actionability without brittle post hoc parsing, each call is constrained to return exactly one JSON object whose required fields, types, and enumerated values are enforced at generation time using Structured Outputs with a strict JSON Schema (“json_schema”); this guarantees structural validity (such as no missing keys or invalid scoring values), while the accompanying evidence ledger preserves transparency for scientific review.

### Implementations in LLMs

We submitted all LLM queries through the OpenAI Python API library (v2.14.0) using a standardized inference interface (client.responses.create), maintaining a constant request structure across models while varying only the model identifier. Each request comprised a fixed developer instruction containing our domain-grounded rubric and a per-pair user message specifying the target gene and indication. We enforced structured output by supplying a JSON schema via text.format with type=“json_schema”. Each T-I pair was queried once, and all experiments were conducted in November 2025.

We evaluated three OpenAI models: GPT-5 (gpt-5-2025-08-07), GPT-4o (gpt-4o-2024-08-06), and GPT-3.5 Turbo (gpt-3.5-turbo-0125). For GPT-5, we disabled tool use (such as web search and function calling) so that generation relied solely on the model’s internal knowledge, ensuring that the evaluation reflected model behavior under our fixed protocol and leakage constraints. We set reasoning_effort = “high” and verbosity = “high”, and extended the timeout to 1,800 seconds per request to accommodate the model’s longer reasoning process. Our primary analysis used the Streaming API, which returns responses synchronously in real time. To assess robustness, we also conducted independent runs using the Batch API, which processes multiple requests asynchronously as queued jobs. GPT-4o and GPT-3.5 were also accessed via the Streaming API. Notably, GPT-3.5 Turbo failed to produce valid structured output given our prompt complexity and was excluded from further analysis. A detailed summary of model-specific settings, cost per query, and latency is provided in **Table S1**.

### Metrics and statistical analysis with drug development data

We evaluated whether T-I pairs supported by LLM-derived genetic evidence have a higher probability of clinical development success using a pre-curated drug development program dataset developed by Minikel, et al. ^1^, originally derived from Citeline Pharmaprojects. This database tracks drug development programs from preclinical stages through all clinical phases and eventual market launch. Minikel, et al. ^1^ curated the dataset using several domain-specific principles. The unit of analysis was the T-I pair, defined as a drug target evaluated for a specific disease indication in drugs. Each T-I pair was annotated with the highest stage of clinical development achieved (phase I, phase II, phase III, or launched) across all drugs targeting that gene for the given indication. They restricted analyses to monotherapy programs initiated since 2000 that had a mapped human gene target and an indication represented in the Medical Subject Headings (MeSH) ontology; combination therapies, diagnostic indications, and records lacking either a human target gene or a MeSH-mapped indication were excluded. They further filtered indications to those with genetic insight, defined as diseases or related phenotypes for which human genetic studies have been successfully conducted. This resulted in 13,022 T-I pairs reaching at least phase I, which we used in our analysis, consistent with their original study.

Following previous phase-transition analyses^1^, phase-specific success is defined as a T-I pair transitioning from a reference phase to the subsequent phase, operationalized using the highest phase reached. For example, a pair is “phase I to phase II” success if the pair reached at least phase II. Overall success from phase I to launched was defined as having reached launched status among T-I pairs that reached at least phase I. For each genetic evidence system (either human-curated or LLM-derived), we computed the probability of genetic support, P(G), defined as the proportion of T-I pairs classified as genetically supported. Relative success (RS) was defined as the risk ratio comparing supported to unsupported programmes: if *N_G_* and *N*_!*G*_ denote the number of T-I pairs that reached the reference phase with and without genetic support, and *X_G_* and *X*_!*G*_ denote the subset that achieved the success endpoint, then RS = (*X_G_*/*N_G_*)⁄(*X*_!*G*_/*N*_!*G*_). We used the Katz logarithmic method for confidence intervals of a risk ratio. For LLM-derived genetic support, we dichotomized integrated causal evidence score (ICES) into supported versus unsupported using the threshold ICES ≥ 4.0 (strong or very strong), consistent with the calibrated rubric described above. Channel-level analyses used the analogous thresholding of channel scores when channel-specific support was required.

Therapeutic area stratification was performed using MeSH’s hierarchical organization: each indication’s MeSH term was mapped to one or more top-level therapeutic areas based on its position in the MeSH tree, enabling estimation of P(G) and RS within therapeutic areas. Cross-area concordance between LLM-derived and human-curated metrics was quantified using rank-based correlations (Spearman) when comparing area-level summaries and linear correlations (Pearson) when comparing continuous ICES values between models; exact statistical tests for each figure are reported in the corresponding figure legends. All statistical analyses were conducted in R 4.3.3.

### Benchmarking with human expert-curated genetic evidence

We benchmarked LLM-derived genetic support against a recent large-scale, human expert-curated integration of genetic evidence reported by Minikel, et al. ^1^. In their framework, genetic support for a T-I pair is defined by linking a target gene to a human genetic association for a trait whose MeSH term is similar to the indication MeSH term (similarity threshold ≥ 0.8). Their analysis primarily focused on the above-mentioned clinical success benchmark universe comprising 13,022 T-I pairs that reached at least Phase I. The curated genetic associations were assembled from multiple complementary resources spanning both Mendelian and complex trait evidence, including OMIM^43^ (https://www.omim.org/) for Mendelian gene-disease associations, Open Targets^8^ (https://www.opentargets.org/) for genetic associations with locus-to-gene prioritization, colocalization resources such as PICCOLO^44^, exome sequencing resources such as Genebass^45^ (https://app.genebass.org/), and somatic mutation resources such as IntOGen^46^ (https://www.intogen.org/). Notably, mapping indication names to MeSH terms, computing semantic similarity between MeSH terms, and integrating genetic evidence across these heterogeneous resources require substantial domain expertise and are nontrivial processes that typically take experts months or even years to complete. In contrast, our LLM-derived framework operates directly on the indication name and does not require these time-consuming manual curation steps.

For exact reproducibility, we used the publicly available code released by Minikel, et al. ^1^. We first verified that, using the 13,022 T-I pairs and the processed genetic-evidence tables, we could reproduce the published figures and summary statistics (P(G) and RS). We then compared statistics computed from human expert-curated genetic-evidence labels with those computed from LLM-derived labels obtained by thresholding ICES, both overall and stratified by therapeutic area.

### TigerAI platform design and implementation

TigerAI platform is a cloud-based platform hosted on AWS that provides researchers with AI-powered genetic evidence analysis for T-I pairs. The platform is built on Django 5.0 (https://www.djangoproject.com/) with GraphQL via Graphene-Django (https://docs.graphene-python.org/projects/django/) for flexible data querying. Data storage employs a multi-database architecture: SQLite (https://sqlite.org/) handles user data and session management, while pre-computed datasets are stored in Apache Parquet format (https://parquet.apache.org/) for efficient columnar access. Static databases contain the T-I universe and gene name and MeSH term mappings, and dynamic storage organizes user-generated analyses by username to support multi-user environments. A PostgreSQL backbone has been implemented to facilitate future scalability as user demand grows.

The public TigerAI interface supports two primary workflows. The AI-generation workflow (https://tigerai.bio/) is our main user-facing service. Users specify a gene target and a disease indication (supported by autocomplete search), which triggers a live LLM call to produce an on-demand, AI-powered genetic support report for that T-I pair. The Browse workflow (https://tigerai.bio/browse), complemented by a benchmarking dashboard, provides the access to pre-computed results across our benchmark universe of 13,022 labeled T-I pairs. In Browse, users can filter by development phase, inspect model-specific verdicts, apply ICES-based filtering, and compare LLM outputs with human expert-curated results from Minikel, et al. ^1^. We also provide benchmarking summaries based on the same labeled dataset (https://tigerai.bio/benchmark).

## Code and data availability

Our platform can be accessed at https://tigerai.bio/. The drug development and human expert-curated genetic evidence data can be obtained from https://github.com/ericminikel/genetic_support/. All data generated by the present study can be found on our platform https://tigerai.bio/.

