## Supplementary material for "TigerAI: An AI-powered genetic evidence platform to support clinical development": supp_information

**This PDF file includes:**

Supplementary Figures S1-S5

**Other Supplementary Materials for this manuscript include the following:**

Supplementary Table S1 (.xlsx) (available in an Excel file)

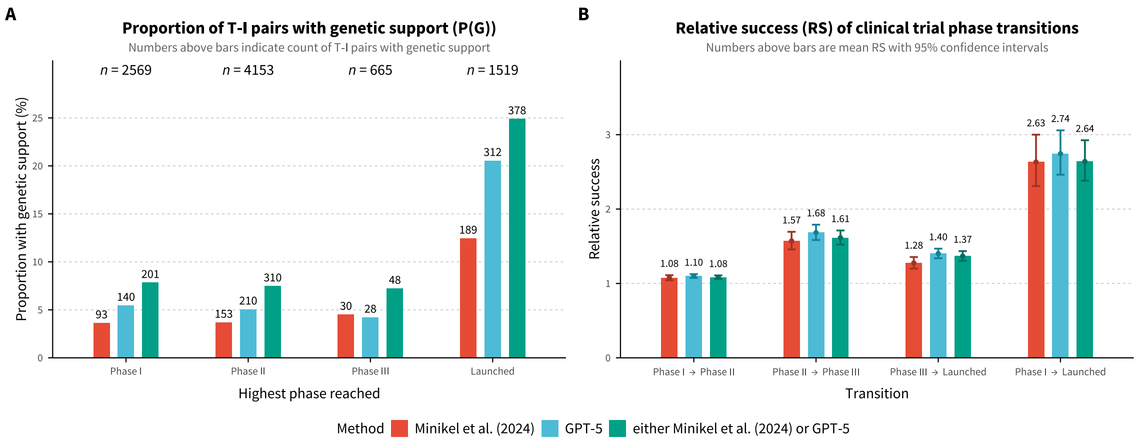


**Fig. S1 Genetic support and clinical trial success using the union of human curation and GPT-5.** (**A**) Proportion of target-indication (T-I) pairs with genetic support, P(G), stratified by highest clinical phase reached and classification method. T-I pairs were classified as genetically supported by human curation using Minikel et al. (2024) alone, GPT-5 alone, or either method (union). Numbers above bars indicate counts of T-I pairs with genetic support. (**B**) Relative success (RS) of clinical trial phase transitions, defined as the ratio of transition probability for genetically supported T-I pairs to that of unsupported pairs, stratified by classification method. Numbers above bars indicate mean RS values; error bars represent 95% confidence intervals calculated using the Katz log method. Analyses use 13,022 T-I pairs from monotherapy programmes added since 2000 that reached at least Phase I; GPT-5 evidence was generated using the domain-grounded instruction described in this study.

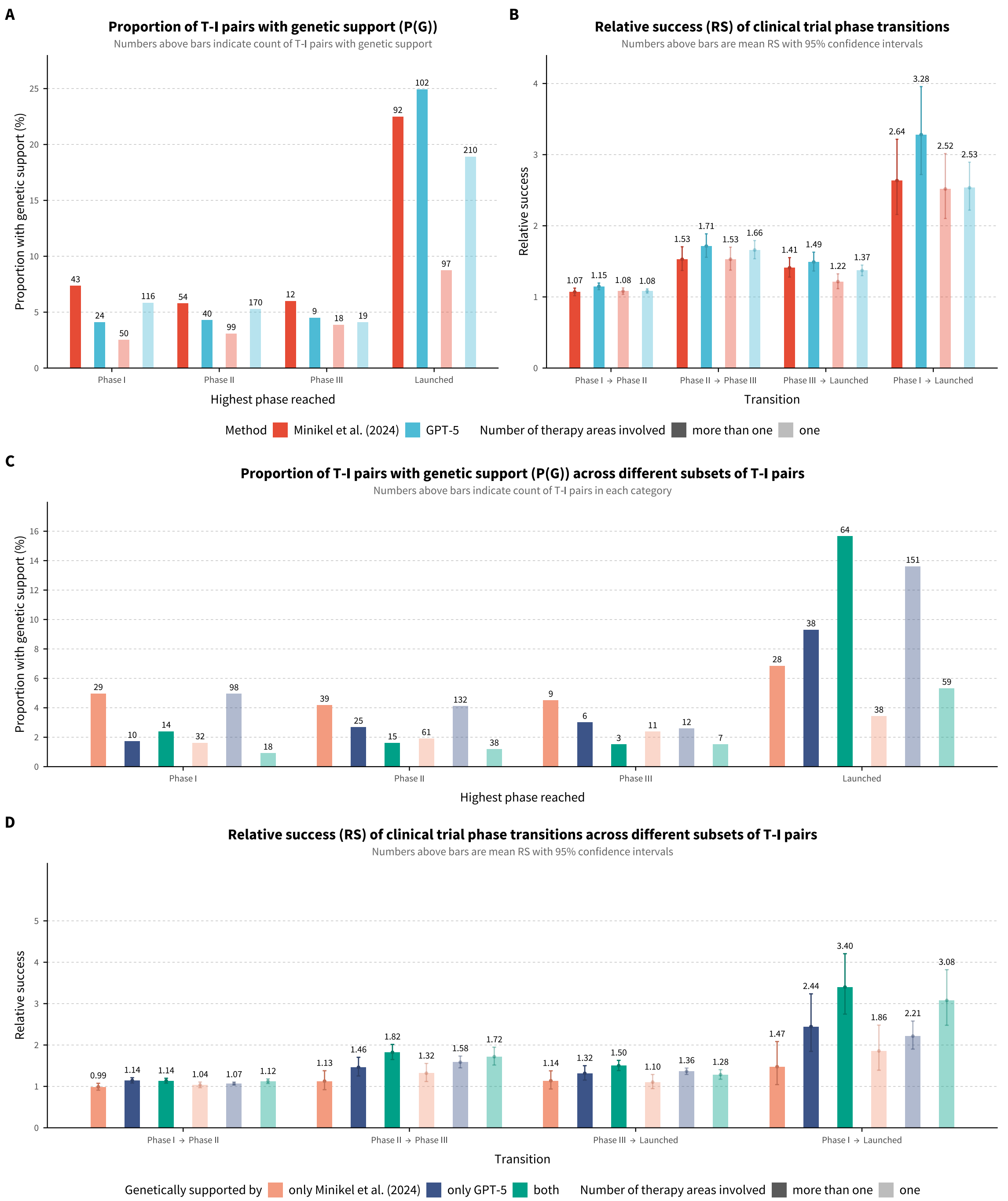


**Fig. S2 Comparison of genetic support metrics between human curation and GPT-5 stratified by the number of therapy areas involved.** (**A**) Proportion of target-indication (T-I) pairs with genetic support, P(G), stratified by number of therapy areas (more than one vs. one) and highest clinical phase reached. Red bars indicate human curation by Minikel et al. (2024); blue bars indicate GPT-5.

Darker shades represent T-I pairs involving more than one therapy area; lighter shades represent T-I pairs involving a single therapy area. Numbers above bars indicate counts of genetically supported T-I pairs. (**B**) Relative success (RS) of clinical trial phase transitions, comparing methods and number of therapy areas involved. RS is defined as the ratio of transition probability for genetically supported T-I pairs to that of unsupported pairs. Points indicate mean RS values; error bars represent 95% confidence intervals calculated using the Katz log method. (**C**) Proportion of T-I pairs with genetic support across mutually exclusive subsets: supported only by human curation (coral), only by GPT-5 (dark blue), or by both methods (teal). Numbers above bars indicate counts of T-I pairs in each category. (**D**) Relative success of clinical trial phase transitions across the same mutually exclusive subsets. T-I pairs supported by both methods show the highest RS for the Phase I to Launched transition (RS = 3.40 for more than one therapy area; RS = 3.08 for one therapy area).

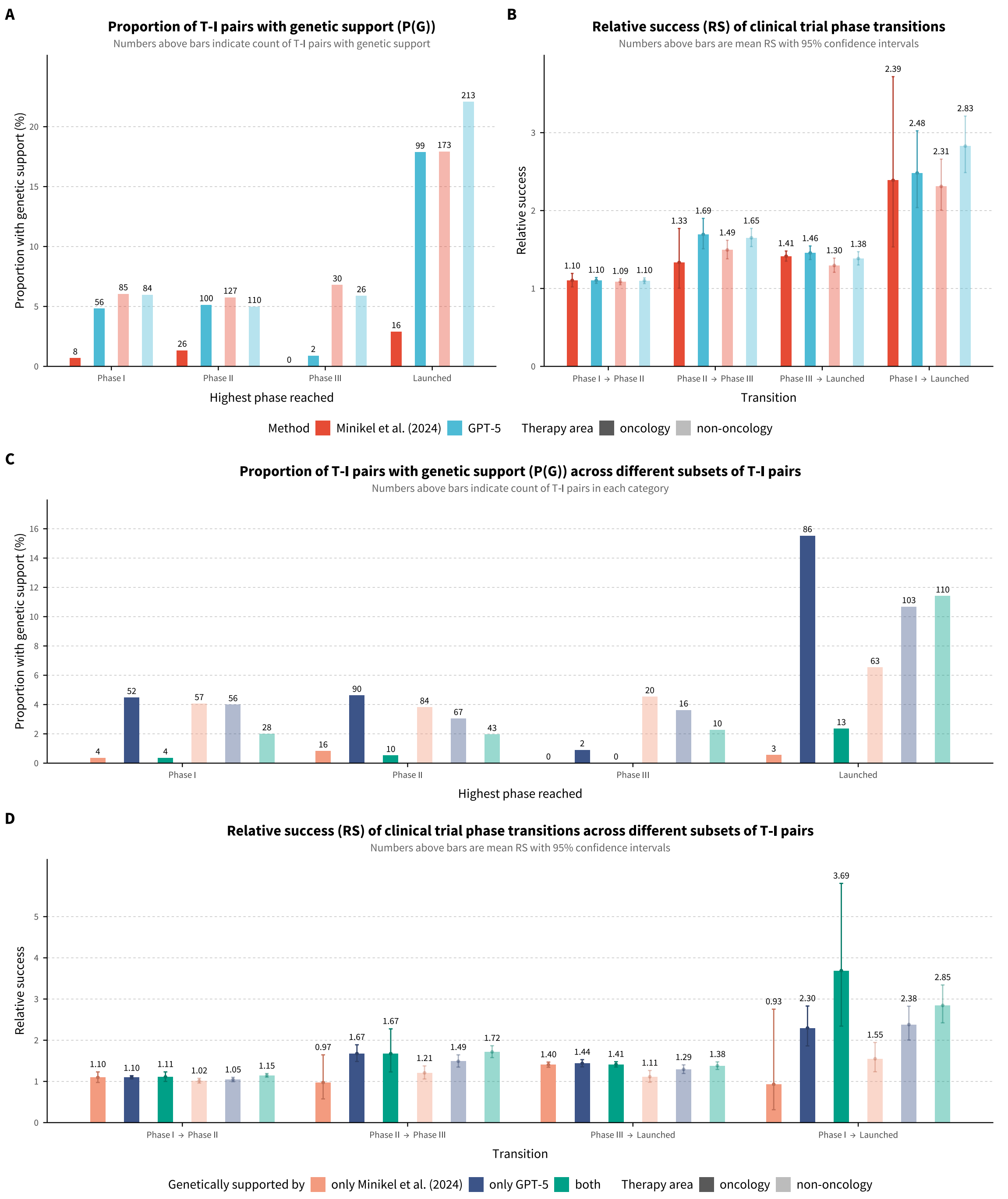


**Fig. S3 Comparison of genetic support metrics between human curation and GPT-5 stratified by whether therapy area involves oncology.** (**A**) Proportion of target-indication (T-I) pairs with genetic support, P(G), stratified by number of therapy context (oncology vs. non-oncology) and highest clinical phase reached. Red bars indicate human curation by Minikel et al. (2024); blue bars indicate GPT-

5. Darker shades represent oncology; lighter shades represent non-oncology. Numbers above bars indicate counts of genetically supported T-I pairs. (**B**) Relative success (RS) of clinical trial phase transitions, comparing methods and therapy context (oncology vs. non-oncology). RS is defined as the ratio of transition probability for genetically supported T-I pairs to that of unsupported pairs. Points indicate mean RS values; error bars represent 95% confidence intervals calculated using the Katz log method. (**C**) Proportion of T-I pairs with genetic support across mutually exclusive subsets: supported only by human curation (coral), only by GPT-5 (dark blue), or by both methods (teal). Numbers above bars indicate counts of T-I pairs in each category. (**D**) Relative success of clinical trial phase transitions across the same mutually exclusive subsets. T-I pairs supported by both methods show the highest RS for the Phase I to Launched transition (RS = 3.69 for oncology; RS = 2.85 for non-oncology).

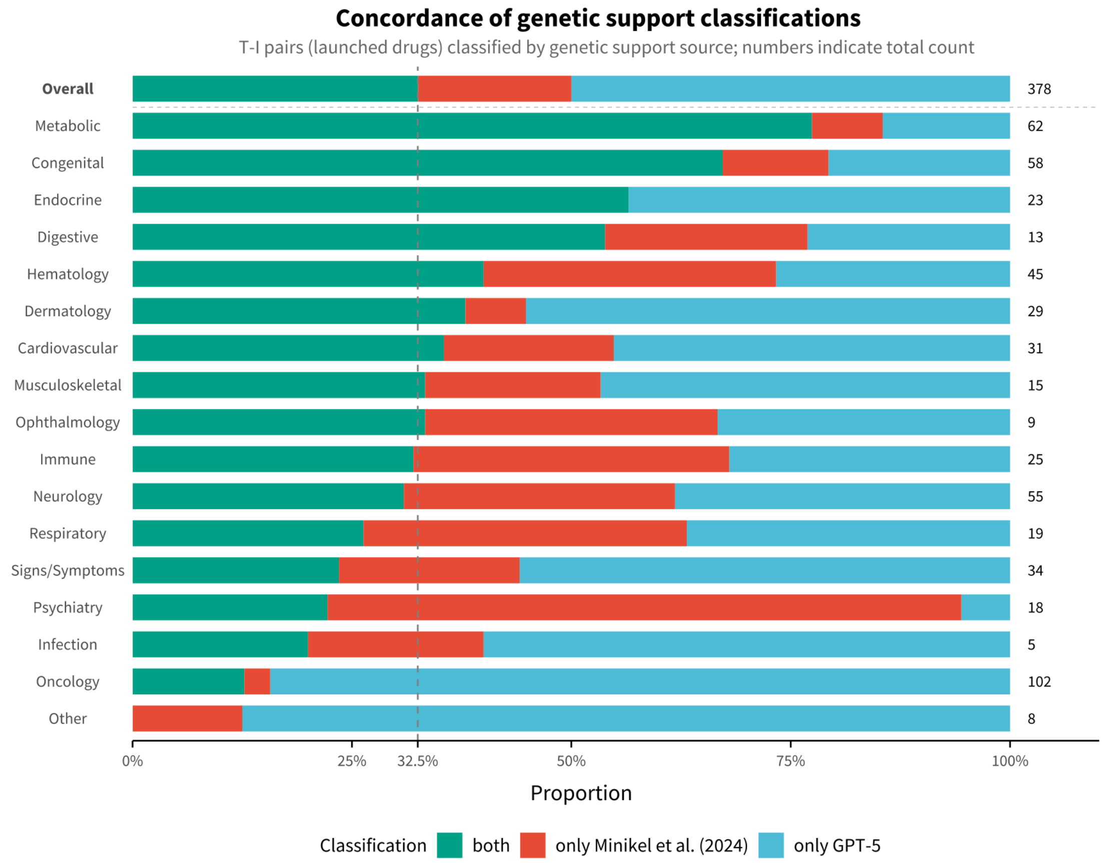


**Fig. S4 Concordance of genetic support classifications between human curation and GPT-5 across therapeutic areas.** Horizontal stacked bar chart showing the proportion of target-indication (T-I) pairs with genetic support, stratified by source of support: both methods (teal), only Minikel et al. (2024) human curation (coral), or only GPT-5 (light blue). Analysis is restricted to T-I pairs corresponding to launched programmes that received genetic support from at least one method. The "Overall" category (top, bold) summarizes all 378 T-I pairs meeting inclusion criteria. Therapy areas are ordered by decreasing proportion of concordant support (both methods). Numbers to the right of each bar indicate the total count of genetically supported T-I pairs in that category. The vertical dashed line at 32.5% marks the overall proportion of T-I pairs supported by both methods. T-I pairs may contribute to multiple therapeutic areas if the indication spans more than one disease category.

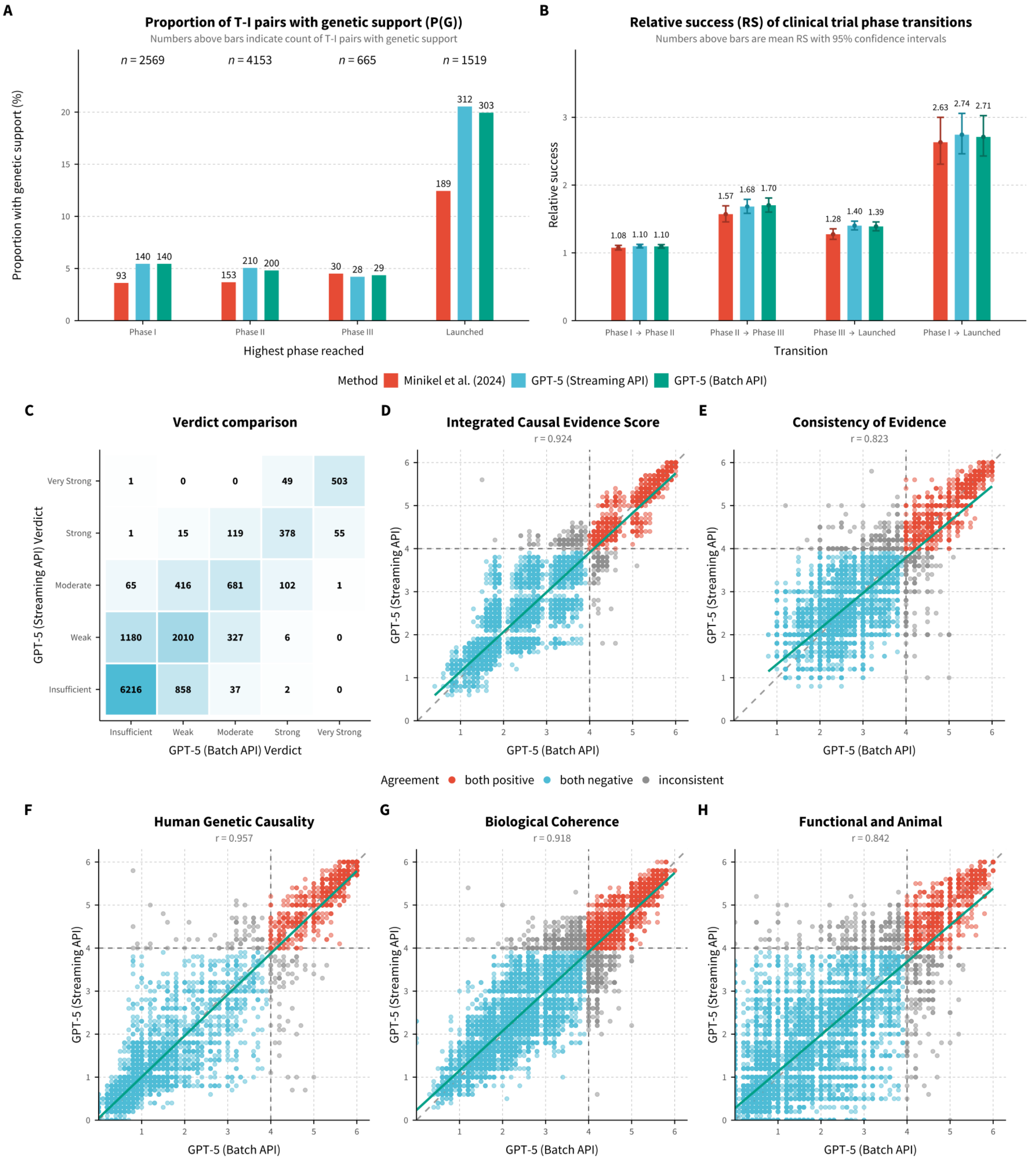


**Fig. S5 Consistency of GPT-5 genetic evidence classification across Streaming and Batch APIs.** (**A**) Proportion of target-indication (T-I) pairs with genetic support, P(G), stratified by highest clinical phase reached and classification method. T-I pairs were classified using human curation by Minikel et al. (2024), GPT-5 Batch API, or GPT-5 Streaming API. Numbers above bars indicate counts of T-I pairs with genetic support. (**B**) Relative success (RS) of clinical trial phase transitions, stratified by classification method. Numbers above bars indicate mean RS values; error bars represent 95% confidence intervals. (**C**) Confusion matrix comparing verdict labels assigned by GPT-5 Batch API and GPT-5 Streaming API, which demonstrates high concordance across independent API calls. Cell values indicate counts of T–I pairs receiving each verdict combination. (**D-H**) Agreement of score components between GPT-5 Batch API and GPT-5 Streaming API, shown as scatterplots for the integrated causal evidence score (**D**) and the four evidence dimensions: consistency (**E**), human genetic causality (**F**), biological coherence (**G**), and functional/animal support (**H**). High Pearson correlation coefficients (r) are reported across all metrics, indicating strong reproducibility of GPT-5 across different processing batches. Points are colored by verdict agreement (both supported: strong/very strong; both unsupported: insufficient/weak/moderate; or discordant), dashed diagonals indicate perfect agreement, and solid lines show linear regression fits.
